# Linear ubiquitin chain assembly complex contributes to NLRP3-mediated pyroptotic cell death

**DOI:** 10.1101/2025.11.18.689012

**Authors:** Tiphaine Douanne, Rosalie Moreau, Valeria Trapani, Kilian Trillet, Hubert Leloup, Virginie Petrilli, Julie Gavard, Nicolas Bidère

## Abstract

Activation of the NLRP3 inflammasome by infectious or sterile insults culminates in pyroptosis, a lytic and highly inflammatory form of programmed cell death. A safeguarded two-step process tightly regulates pyroptosis: priming, which drives NF-κB signaling, followed by execution, ultimately leading to plasma membrane rupture. Linear (Met1-linked) ubiquitination, catalyzed by the E3 ligase complex LUBAC, was previously shown to participate in pyroptosis, but the underlying mechanisms are not fully understood. In this study, we show that Met1-linked ubiquitin chains can assemble during both priming and execution phases, independently of the inflammasome sensor NLRP3. Genetic deletion of the LUBAC enzymes or pharmacological inhibition impaired pyroptosis. Conversely, cell death was enhanced without the deubiquitinase OTULIN, which selectively removes linear ubiquitination. Finally, using an optogenetic model to bypass priming, we demonstrate that Met-1-linked ubiquitination is required for the execution phase of pyroptosis. These findings offer insights into the regulation of pyroptotic cell death by linear ubiquitination.

## INTRODUCTION

Programmed cell death shapes human life, with multiple mechanisms coexisting to preserve tissue homeostasis and regulate inflammatory responses ^1^. Among these, pyroptosis is a lytic and highly inflammatory form of cell death that plays a central role in host defense against infection. A variety of sensors, named inflammasomes, assemble in response to diverse pathogenic or danger signals to activate caspase-1 (CASP1), with the NLRP3 inflammasome amongst the most broadly responsive. NLRP3-mediated pyroptosis is a finely tuned, two-step process that ensures the inflammatory outcome occurs only under conditions of genuine pathogenic threat. The first signal, or priming, is typically initiated by pathogen-associated molecular patterns (PAMPs), such as the bacterial wall component lipopolysaccharide (LPS), or by certain danger-associated molecular patterns (DAMPs). These molecules activate pattern recognition receptors (PRRs), including Toll-like receptors (TLRs), leading to activation of the NF-κB pathway and transcriptional upregulation of key components of the pyroptotic machinery, such as *NLRP3, pro-IL-1*β, and *pro-IL-18* ^2^. However, priming alone is not sufficient to trigger NLRP3-mediated pyroptosis, and a second signal, often another PAMP or DAMP, such as crystalline substances, pore-forming toxins, or nucleic acids, is required to activate the execution phase ^3^. For example, the pore-forming toxin Nigericin acts as a potassium ionophore and drives the formation of a supramolecular complex scaffolded by NLRP3, known as the inflammasome ^4^. Upon sensing activating stimuli, NLRP3 recruits the adaptor protein ASC (apoptosis-associated speck-like protein containing a CARD), which undergoes prion-like polymerization to form large ASC specks ^5^. ASC, in turn, recruits pro–caspase-1 via CARD–CARD interactions, enabling its proximity-induced autoactivation ^6,7^. Active CASP1 then maturates the pro-forms of the inflammatory cytokines IL-1β and IL-18, as well as the pore-forming effector GSDMD. The liberated N-terminal fragment of GSDMD oligomerizes within the plasma membrane to form pores that allow the release of mature IL-1β, IL-18, and DAMPs ^8^. The progressive accumulation of these pores drives ionic imbalance and osmotic swelling, ultimately leading to plasma membrane rupture via NINJ1 oligomerization, release of cytosolic contents, including larger DAMPs such as HMGB1 ^9^, and cell demise ^10^.

The activity of inflammasome components is precisely regulated by post-translational modifications (PTMs), which control their assembly and signaling output. Among these, phosphorylation and ubiquitination critically modulate NLRP3 activation ^11,12^. Ubiquitination consists of the covalent attachment of ubiquitin to lysine residues on substrates as single units or polyubiquitin chains with distinct linkages, governing protein stability, activity, localization, and signaling, thereby regulating most cellular processes in eukaryotic cells ^13^. This PTM is reversible through the action of deubiquitinating enzymes (DUBs), a group of proteases that selectively remove ubiquitin from target proteins. Among the linkage types, linear (“head-to-tail” or Met1-linked) ubiquitination is unique, as it is exclusively catalyzed by the Linear Ubiquitin Chain Assembly Complex (LUBAC), which comprises the E3 ligases HOIP, HOIL-1, and the adaptor SHARPIN. Along this unique enzymatic complex, only two DUBs, OTULIN and CYLD, are known to target linear ubiquitin chains. OTULIN is specific for Met1 linkages, whereas CYLD hydrolyzes both Met1- and Lys63-linked ubiquitin chains ^14,15^. CYLD has been shown to restrain pyroptosis ^16,17^. The LUBAC has emerged as an essential gateway for NF-κB activation downstream numerous immunoreceptors, including PRRs, controlling key genes involved in inflammation and cell survival ^18^. In addition, the LUBAC has also been linked with pyroptosis. Mice deficient for SHARPIN (also known as *Cpdm*) develop chronic proliferative dermatitis, and this phenotype is attenuated or delayed by genetic deletions of *Nlrp3* ^19^, *Casp1* ^19^, and *Il1r* ^20,21^. Moreover, SHARPIN was shown to bind and limit CASP1 activity, inhibiting maturation of IL-1β and IL-18. Notably, this function appears to be independent of LUBAC activity ^22^. An early study using bone marrow-derived macrophages from *Hoil-1*-deficient mice described a role for the E3 ligase in ASC linear ubiquitination and NLRP3 inflammasome assembly ^23^. More recently, a pathogenic phosphorylation of HOIL-1 was shown to exclude it from LUBAC, thereby impairing linear ubiquitination of ASC and inhibiting NLRP3 inflammasome assembly. In parallel, this phosphorylation stabilizes and activates HOIL-1, promoting Lys48-linked ubiquitination and proteasomal degradation of NLRP3 ^24^. However, despite the emerging connection between LUBAC components and inflammasome regulation, the specific contribution of linear ubiquitination and the deubiquitinase OTULIN to the execution remains poorly understood.

In this study, using a combination of genetic deletions, chemical inhibition, and an optogenetic model, we demonstrate that LUBAC-mediated linear ubiquitination operates at multiple steps of the NLRP3-mediated pyroptotic process to positively regulate cell death, notably during its execution phase. We also show that cells lacking OTULIN display enhanced pyroptotic cell death, further underscoring the pro-pyroptotic functions of linear ubiquitination.

## RESULTS

### The execution step of pyroptosis drives linear ubiquitination

To induce pyroptotic cell death, THP-1 human monocytic cells were subjected to a two-step stimulation: cells were first primed with LPS and subsequently treated with Nigericin (**Figure 1A**). Upon exposure to Nigericin, cells underwent pyroptosis, as evidenced by GSDMD cleavage and the release of the DAMP HMGB1 into the culture supernatant, and this response was efficiently suppressed by treatment with the NLRP3 inhibitor MCC-950 (**Figure 1B**). Consistent with a rupture of the plasma membrane, we found that all LUBAC subunits, together with the associated deubiquitinase OTULIN, were released into the supernatant upon pyroptotic cell death (**Figure 1C**). In addition, the execution phase of pyroptosis was accompanied by a change in the pattern of linear ubiquitination, with a portion of Met1-linked ubiquitin chains detected in the culture supernatant (**Figure 1B**). Moreover, lysates from cells treated with MCC-950 showed increased levels of linear ubiquitin chains, suggesting an induction upstream of NLRP3 activation (**Figure 1B**). By contrast, Lys63-linked ubiquitination dynamics remained essentially unchanged (**Figure 1B**). Further supporting an induction of linear ubiquitination independent of NLRP3, Nigericin treatment alone also enhanced Met1-ubiquitination in A549 cells, which lack NLRP3 expression (**Figure 1D**). Because a fraction of linear ubiquitin chains appeared in the supernatants of pyroptotic cells, we next used HMGB1-Lucia-expressing THP-1 cells, deleted for GSDMD (GSDMD^sg^), in which pore formation and subsequent cell lysis and HMGB1 release are prevented (**Figure 1E and 1F**). Importantly, these cells exhibited a marked accumulation of Met1-linked ubiquitination was observed following LPS/Nigericin stimulation, confirming the formation of linear ubiquitin chains during the execution of pyroptosis (**Figure 1E**). Thus, linear ubiquitination dynamics are changed independently of NLRP3 during the execution phase of pyroptosis.

**Figure 1.**
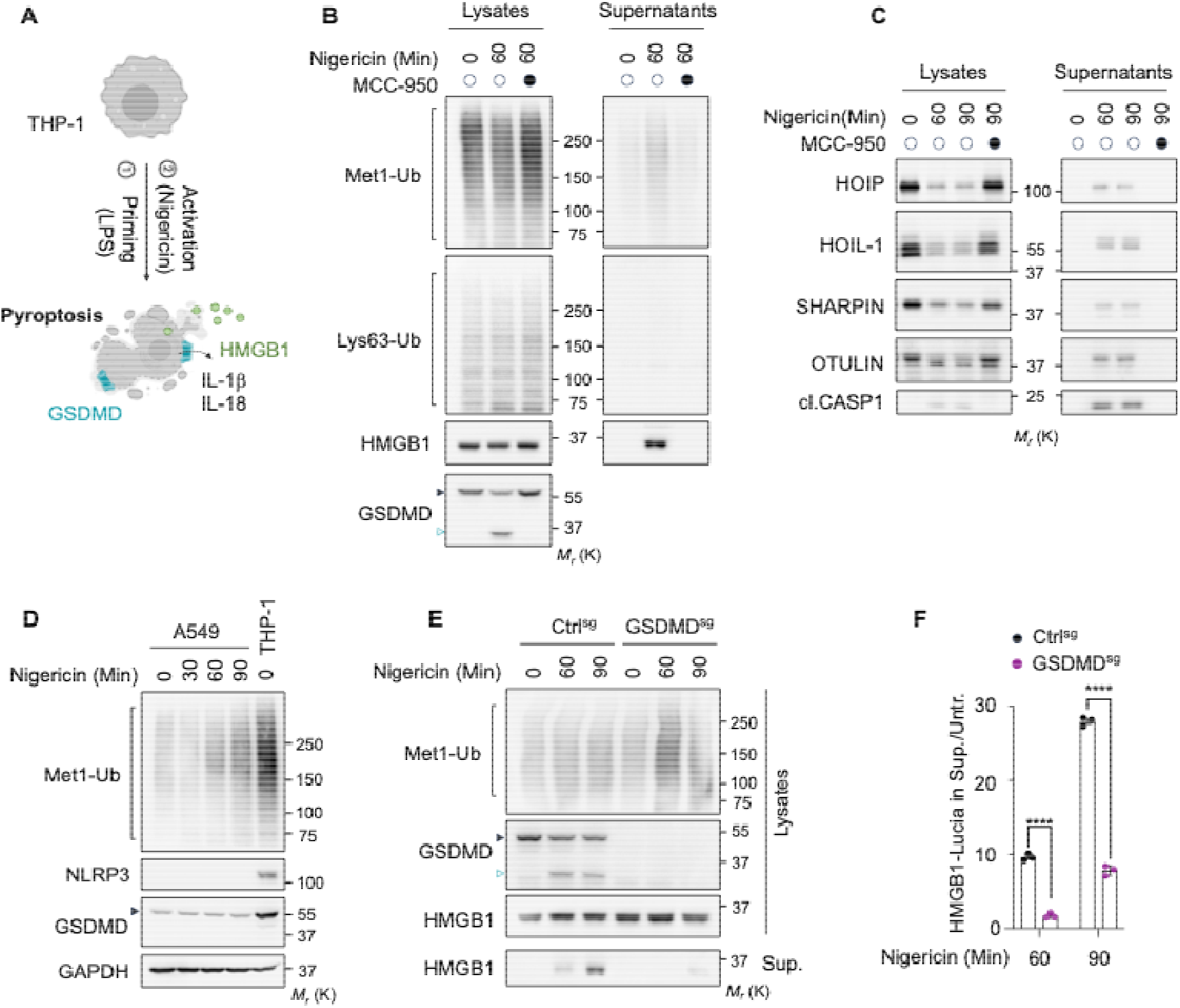
Pyroptosis execution drives the formation of linear ubiquitin chains. (**A**) Workflow used to induce pyroptosis in THP-1 monocytic cells. (**B**) Immunoblots of indicated proteins in lysates and supernatants of THP-1 cells primed with 1 µg/mL of LPS for 3 hours before treatment with 10 µM of Nigericin as indicated. Where indicated, cells were pre-treated with 1 µM of MCC-950 for 30 Min before Nigericin addition. Arrowheads indicate protein-cleaved fragments. The positions of molecular weights (Mr) are indicated. (**C**) Immunoblots of THP-1 treated and analyzed as in (B). **(D)** Immunoblots of indicated proteins in lysates of A549 cells treated as indicated. Lysates from untreated THP-1 cells serve as a control. (**E**) THP-1 cells stably expressing HMGB1-Lucia deleted with sgRNAs for GSDMD (GSDMD^sg^), or their control counterparts (Ctrl^sg^) were treated and analyzed as in (B). (**F**) The release of HMGB1-Lucia was assessed in the supernatants of cells as in (E). (n=3; mean ± SEM; ^****^P<0.0001; t-test). All the presented data are representative of at least three independent experiments.

### The LUBAC is required for the efficient execution of pyroptosis

We next examined the impact of inhibiting linear ubiquitin chain formation on pyroptosis. To this end, we generated THP-1 cells lacking both E3 ligases HOIP and HOIL-1 using specific single guide RNAs (HOIP^sg^ and HOIL-1^sg^). As expected, the deletion of HOIP and HOIL-1 markedly impaired the formation of Met1-linked ubiquitin chains (**Figure 2A-C**). Moreover, the loss of HOIP and HOIL-1 strongly reduced GSDMD processing and diminished HMGB1 release into the supernatant (**Figure 2A-C**). Consistently, plasma membrane rupture, monitored by Sytox Green incorporation, was significantly attenuated in the absence of HOIP and HOIL-1 (**Figure 2D**). To next assess if the catalytic activity of LUBAC is required during pyroptosis, HMGB1-Lucia THP-1 cells were pretreated with the small-molecule selective HOIP inhibitors HOIPIN-1 and HOIPIN-8^25^ following LPS priming and before Nigericin stimulation. HOIPIN treatment efficiently reduced Nigericin-mediated linear ubiquitin chain formation, GSDMD processing, HMGB1 release into the supernatant, and plasma membrane rupture (**Figure 2E and 2G**). Together, these findings demonstrate that LUBAC catalytic activity is essential for efficient execution of pyroptosis, supporting a role for linear ubiquitination in this form of cell death.

**Figure 2.**
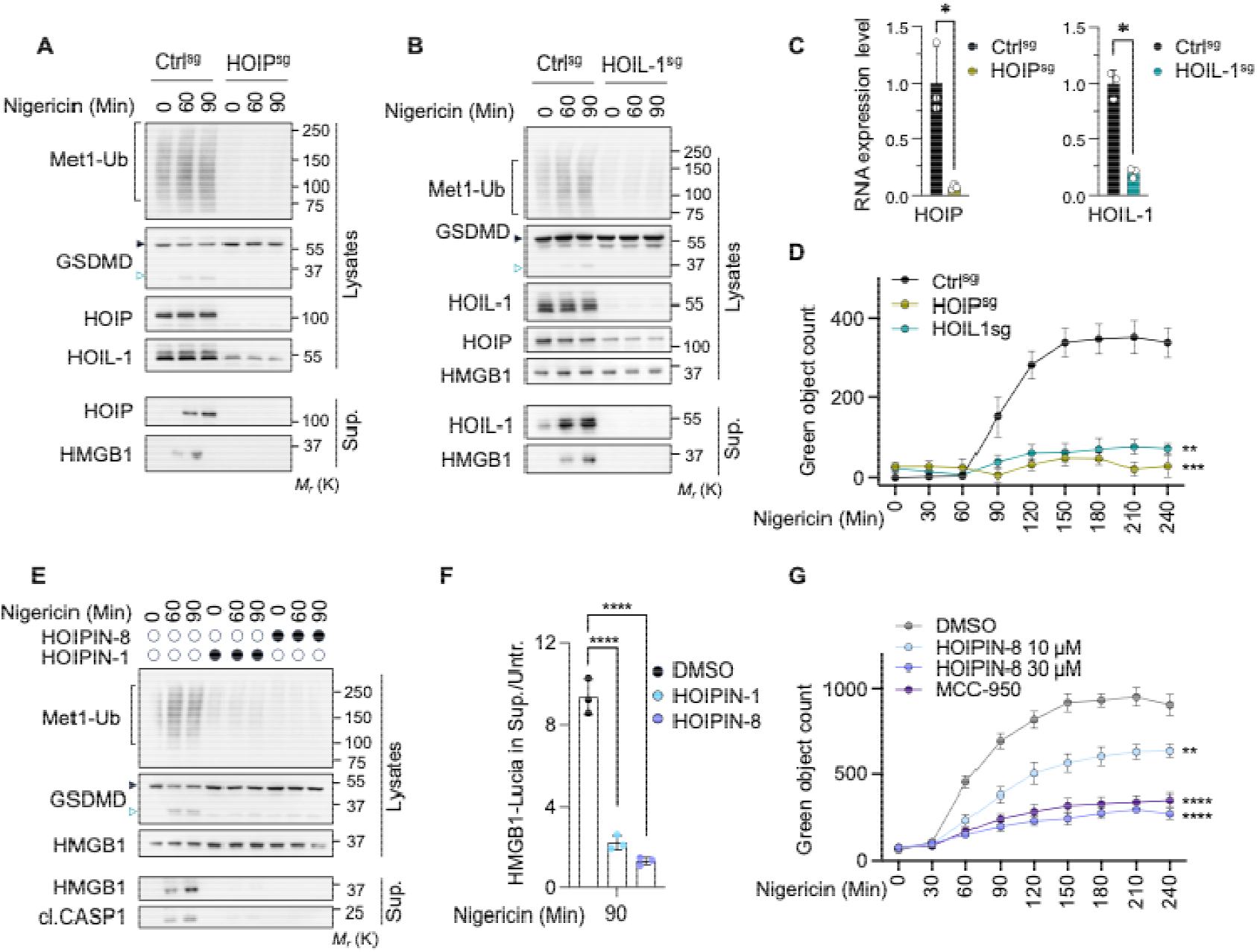
LUBAC and linear ubiquitination drive pyroptosis. (**A**) Immunoblots of indicated proteins in lysates and supernatants (Sup.) of THP-1 cells deleted with sgRNAs for HOIP (HOIP^sg^), or their control counterparts (Ctrl^sg^), primed with 1 µg/mL of LPS for 3 hours before treatment with 10 µM of Nigericin as indicated. Arrowheads indicate protein-cleaved fragments. The positions of molecular weights (Mr) are indicated. (**B**) THP-1 deleted with sgRNAs for HOIL-1 (HOIL-1^sg^), or their control counterparts (Ctrl^sg^), were treated and analyzed as in (A). (**C**) qPCR analysis of HOIP and HOIL-1 in the indicated THP-1 cell genotypes (n=3; mean ± SEM; ^*^P<0.05; t-test). (**D**) PMA-differentiated THP-1 Ctrl^sg^, HOIP^sg^, and HOIL-1^sg^ were stimulated with 10 µM of Nigericin in the presence of 100 nM of Sytox Green. IncuCyte acquisition was performed every 30 Min to assess green object count (n=3; mean ± SEM; ^**^P<0.01, ^***^P<0.001 at end time point; ANOVA). (**E**) THP-1 were pre-treated with 30 µM of HOIPIN-1 or HOIPIN-8, then treated and analyzed as in (A). (**F**) THP-1 cells stably expressing HMGB1-Lucia were treated as in (E). The release of HMGB1-Lucia was assessed in cell supernatants. (n=3; mean ± SEM; ^****^P<0.0001; ANOVA). (**G**) PMA-differentiated THP-1 were pre-treated with 30 µM of HOIPIN-1, HOIPIN-8, or with 1 µM of MCC-950, then treated with 10 µM of Nigericin in the presence of 100 nM of Sytox Green. IncuCyte acquisition was performed as in (D) (n=3; mean ± SEM; ^**^P<0.01, ^****^P<0.0001 at end time point; ANOVA). The presented data are representative of at least three independent experiments.

### OTULIN counteracts pyroptotic cell death

Given our identification of linear ubiquitin as a regulator of pyroptosis, we next examined cell death in cells lacking OTULIN (OTULIN^sg^). Supporting previous reports ^26,27^, deletion of OTULIN led to a pronounced accumulation of overall Met1-linked ubiquitination and enhanced self-ubiquitination of both HOIP and HOIL-1 (**Figure 3A and 3B**). Moreover, OTULIN^sg^ THP-1 cells displayed increased GSDMD processing and elevated release of HMGB1 into the culture supernatant (**Figure 3A and B**), along with enhanced plasma membrane rupture (**Figure 3C**). Together, these findings indicate that loss of OTULIN enhances pyroptotic cell death, establishing OTULIN as a negative regulator of pyroptosis and underscoring the promoting role of linear ubiquitination in pyroptosis.

**Figure 3.**
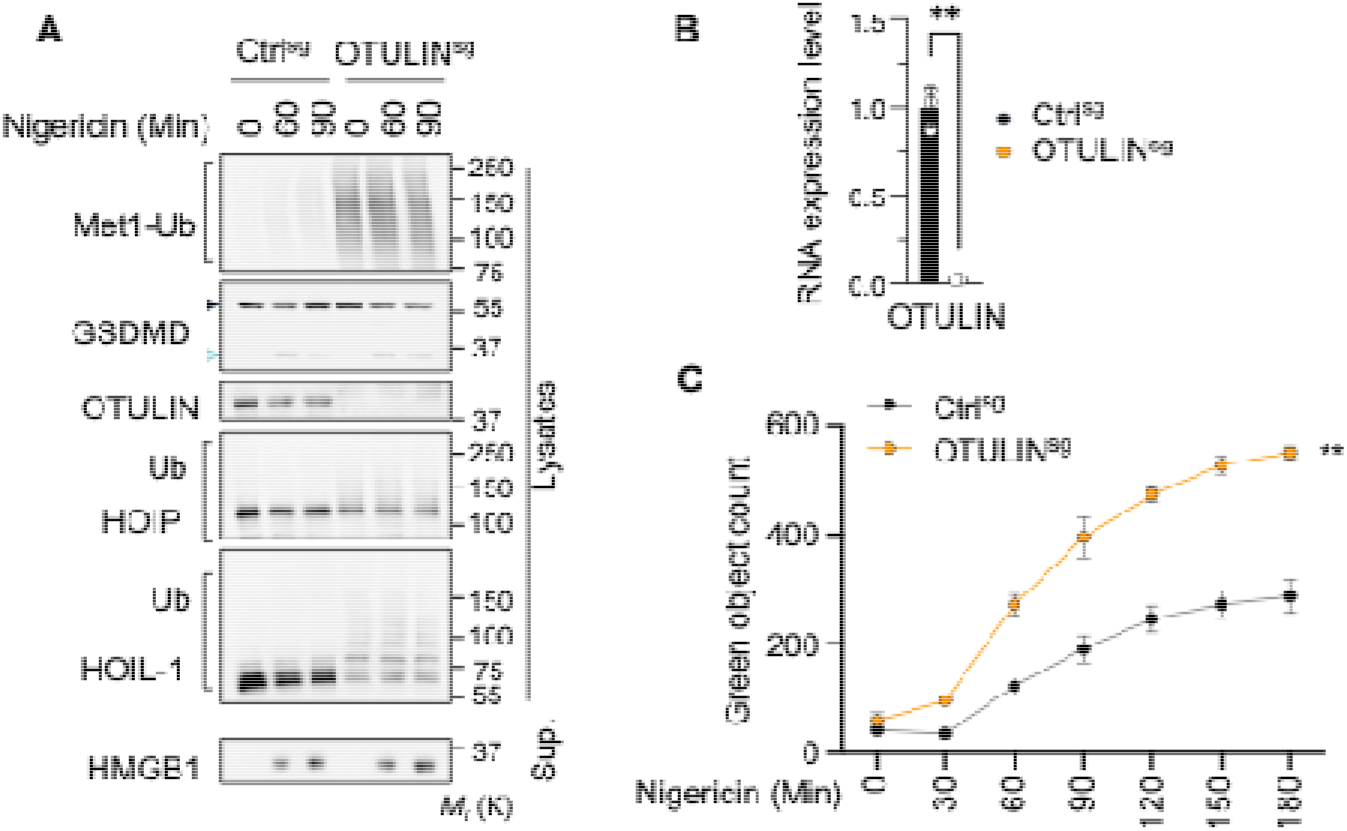
OTULIN counteracts pyroptosis. (**A**) Immunoblots of indicated proteins in lysates and supernatants (Sup.) of THP-1 cells deleted with sgRNAs for OTULIN (OTULIN^sg^), or their control counterparts (Ctrl^sg^), primed with 1µg/mL of LPS for 3 hours before treatment with 10 µM of Nigericin as indicated. Arrowheads indicate protein-cleaved fragments. The positions of molecular weights (Mr) are indicated. (**B**) qPCR analysis of OTULIN in THP-1 Ctrl^sg^ and OTULIN^sg^ (n=3; mean ± SEM; ^**^P<0.01 at end time point; t-test). (**C**) PMA-differentiated Ctrl^sg^ and OTULIN^sg^ THP-1 were stimulated with 10 µM of Nigericin in the presence of 100 nM of Sytox Green. IncuCyte acquisition was performed every 30 Min to assess green object count (n=3; mean ± SEM; ^**^P<0.01, at end time point; t-test). The presented data are representative of at least three independent experiments.

### The LUBAC is required for the execution phase of pyroptosis

The priming of cells is a prerequisite to NLRP3-mediated pyroptosis execution, as it activates the transcription of key inflammasome components and pro-inflammatory cytokines. For instance, LPS engages TLR4, triggering a signaling cascade that culminates in NF-κB activation ^2,28^. Given that LUBAC is a major contributor to NF-κB activation in various physiological contexts ^18^, it may affect pyroptosis execution by regulating the priming phase. Accordingly, LPS treatment in THP-1 cells led to the formation of linear ubiquitin chains, which was impaired in HOIP^sg^ cells (**Figure 4A**). To directly test the contribution of the LUBAC in the execution phase of pyroptosis, we took advantage of a recently developed optogenetic model enabling light-induced oligomerization of the inflammasome adaptor ASC ^29^. This allows for driving pyroptosis independently of priming or secondary activating stimuli (**Figure 4B**). Accordingly, blue-light exposure of opto-ASC THP-1 cells induced GSDMD processing and HMGB1 release (**Figure 4C**). As expected, inhibition of CASP1 with VX-765 reduced both GSDMD cleavage and HMGB1 release. Moreover, consistent with a recent report ^30^, treatment with disulfiram (DSF) did not affect GSDMD processing but prevented pore formation and subsequent HMGB1 release (**Figure 4C**), further validating this optogenetic model. In these conditions, we found that pharmacological inhibition of LUBAC with HOIPIN-8 fully abrogated light-induced pyroptosis (**Figure 4C**). Thus, our data support a critical role for linear ubiquitination in the activation phase of pyroptosis, independent of priming.

**Figure 4.**
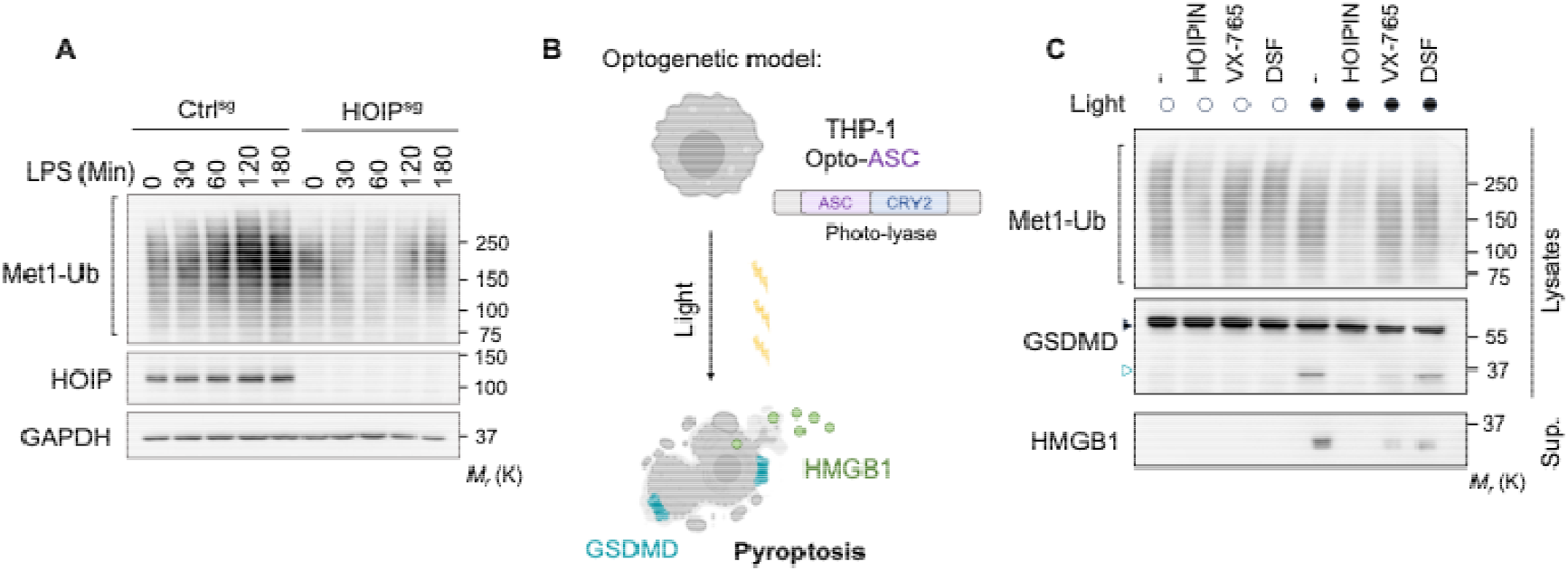
LUBAC and linear ubiquitination contribute to the activation phase of pyroptosis. (**A**) Immunoblots of indicated proteins in lysates of Ctrl^sg^ and HOIP^sg^ THP-1 cells treated with 1 µg/mL of LPS for 30-180 minutes. The positions of molecular weights (Mr) are indicated. Data are representative of at least three independent experiments. (**B**) Optogenetic model to activate pyroptosis independently of priming steps. (**C**) Immunoblots of indicated proteins in lysates and supernatants (Sup.) of opto-ASC THP-1 cells pretreated with 30 µM of HOIPIN-8, 10 µM of VX-765 or 2 µM of disulfiram (DSF) for 30 minutes and exposed to 3 series of 25 x 1s light pulses (477 nm) with an interval of 1 min between each series. The positions of molecular weights (Mr) are indicated. Data are representative of at least three independent experiments.

## DISCUSSION

Pyroptosis is a tightly safeguarded two-step process consisting of priming and activation phases, with priming relying on NF-κB signaling and preconditioning cell death. The LUBAC and linear ubiquitination are known to facilitate NF-κB activation downstream of TLRs, thereby enabling efficient priming. However, the role of the LUBAC in the execution phase of pyroptosis remains incompletely understood. Previous studies showed that linear ubiquitination of the inflammasome component ASC favors NLRP3 assembly and activation ^23,24^. We now report that linear ubiquitin chains also accumulate when NLRP3 is inhibited or in cells lacking NLRP3, suggesting an additional layer of ubiquitin-mediated regulation independent of NLRP3 activation. Whether this accumulation is driven by the K^+^ efflux from the cytosol caused by Nigericin ^4^, and whether other pore-forming toxins act similarly, warrants further investigation. Moreover, additional work will be essential to identify the relevant substrates upstream and downstream of NLRP3 and elucidate their biological significance during pyroptotic cell death. In that view, we used an optogenetic system that drives ASC oligomerization and pyroptosis independently of upstream priming and activation signals, and found that LUBAC is required during the execution phase of pyroptosis. Hence, linear ubiquitination occurs and likely regulates the pyroptotic process at multiple stages.

We also show that the loss of the Met1-ubiquitin chain-specific DUB OTULIN enhances pyroptotic response. This observation parallels the recent description that CYLD, the only other DUB known to remove linear ubiquitin-chains in addition to Lys63-ubiquitin chains, licenses NLPR3 activation ^16^. Whether OTULIN and CYLD compete, cooperate, or act at different levels in the pyroptotic signaling pathway remains to be elucidated. In keeping with this, recent studies have highlighted the importance of branched ubiquitin chains in signaling ^31^, and whether the observed chains are composed solely of Met1-linked moieties or represent mixed or branched architectures incorporating additional ubiquitin linkages, such as Lys63-linked chains, remains to be determined. Nevertheless, our work highlights the importance of dynamic linear ubiquitin turnover in maintaining proper control of pyroptosis.

The identification of LUBAC as a positive regulator of pyroptotic execution expands the functional repertoire of linear ubiquitination beyond its already well-established role in inflammatory signaling. Linear ubiquitin chains are best known for stabilizing signaling complexes downstream of pattern recognition and cytokine receptors, thereby promoting NF-κB activation and cell survival ^18^. Our findings contribute to placing this modification at a later stage of the inflammatory cascade, where it contributes directly to the execution of lytic cell death. Understanding how linear ubiquitination shapes the amplitude and timing of this cell death pathway could reveal new opportunities to therapeutically modulate inflammation in diseases where pyroptosis is detrimental.

## ACKNOWLEDGMENTS

We thank all members of the SOAP laboratory for their help and insightful discussions; Tanguy Deniaud for technical assistance; and the CytoCell facility (SFR Santé François Bonamy, Nantes, France) for the expertise. This work was funded by Ligue Nationale contre le Cancer, Equipe labellisée (JG), Ligue Nationale contre le Cancer, comités de Loire-Atlantique, Maine et Loire, Vendée, Ille et Vilaine, Mayenne, Finistère (NB), Institut National Du Cancer INCa_18384 (NB), Fondation ARC contre le Cancer PJA2021060003886 (NB), SIRIC ILIAD INCA-DGOS-INSERM-ITMO Cancer_18011 (JG). TD was supported by the Institut National du Cancer (INCa). ANR-22-ce15-0032-01 (VP).

## AUTHOR CONTRIBUTIONS

Conceptualization: TD, VP, JG, NB; Methodology: TD, RM, VP, NB; Investigation: TD, RM, VT, KT; Data curation: TD, RM, VT, KT; Visualization: TD, NB; Funding acquisition: JG, NB; Project administration: NB; Supervision: NB; Writing – original draft: TD, NB; Writing – review & editing: TD, KT, VP, JG, NB

## MATERIALS and METHODS

### Cultured Cells

THP-1 and A549 cells were obtained from the American Type Culture Collection (ATCC), and THP-1 cells stably expressing HMGB1-Lucia were purchased from InvivoGen. THP-1 Opto-ASC cells were generated as previously described ^29^. All THP-1 cell lines were cultured in RPMI 1640 medium (Gibco) supplemented with 20% heat-inactivated fetal calf serum (FCS), 1% GlutaMAX (Thermo Fisher Scientific), 1% HEPES (Thermo Fisher Scientific), 1% sodium pyruvate (Thermo Fisher Scientific), 1% MEM non-essential amino acids (Sigma-Aldrich), 1% penicillin-streptomycin (Thermo Fisher Scientific), and 40 ng/mL Normocin (InvivoGen). A549 cells were cultured as THP-1 cells, except the medium was supplemented with 10% FCS.

### Cell death induction and measurement of cell viability

THP-1 cells were washed twice with Opti-MEM (Thermo Fisher Scientific), and 1-2 × 10□ cells were used per condition. Pyroptotic cell death was induced through a two-step stimulation protocol. Cells were first primed with 1 µg/mL lipopolysaccharide (LPS, Invivogen) for 3 hours, followed by stimulation with 10 µM Nigericin for the indicated times (Invivogen). The release of HMGB1 fused to Lucia was measured per the manufacturer’s instructions (Invivogen). In some experiments, THP-1 cells were differentiated/primed using 0.5 µM of PMA (Sigma-Aldrich) for 3 hours, followed by several PBS washes. The next day, cells were treated with 10 µM Nigericin together with 100 nM of Sytox Green (Life Technologies). Green fluorescence was tracked through image acquisition every 30 minutes using the IncuCyte S3 equipment (Sartorius). When indicated, pharmacological inhibitors targeting HOIP (HOIPIN-1 and HOIPIN-8, MedChemExpress, 10-50 µM), NLRP3 (MCC-950, Invivogen, 1 µM), Caspase-1 (VX-765, Selleckchem, 10 µM), or GSDMD (disulfiram or DSF, Santa Cruz, 2 µM) were applied 30 minutes before Nigericin stimulation.

### Lysate and supernatant preparation, immunoblotting

THP-1 cells in suspension were centrifuged at 300*g* for 5 minutes, and the supernatant was collected. Cell pellets were lysed in RIPA buffer [25 mM Tris-HCl (pH 7.4), 150 mM NaCl, 0.1% SDS, 0.5% sodium deoxycholate, 1% NP-40, 1 mM EDTA] supplemented with 1× Halt Protease Inhibitor Cocktail (Thermo Fisher Scientific) for 30 minutes on ice, then cleared by centrifugation at 10,000*g* for 10 minutes. Protein concentration in the lysates was determined using a BCA Protein Assay Kit (UP40840A, Interchim). For immunoblotting, 5–10 µg of total protein from lysates were mixed with 2x Laemmli buffer (Life Technologies) and heated at 95°C for 3 minutes. In parallel, 20 µL of supernatant were mixed with 6× Laemmli buffer (Thermo Fisher Scientific) and boiled under the same conditions. Samples were resolved by SDS-polyacrylamide gel electrophoresis (SDS–PAGE) using 3–15% Tris–acetate gels and transferred onto nitrocellulose membranes (GE Healthcare). The following antibodies were purchased from Santa Cruz: HOIL-1 (H-1, sc-393754), HMGB1 (HAP46.5, sc-56698), GAPDH (6C5, sc-32233). Antibodies to cleaved CASP1 (D57A2, 4199), GSDMD (E9S1X, 39754), OTULIN (14127), and NLRP3 (D4D8T, 1510) were from Cell Signaling Technologies. Antibodies to HOIP (A303-560A) and SHARPIN (A303-559A) were purchased from Bethyl Laboratories. Antibody to linear ubiquitination (1E3, ZRB2114) and Lys63-ubiquitin (clone Apu3) from Sigma were used.

### CRISPR generation and qPCR

Deletion of GSDMD, HOIP, HOIL-1, or OTULIN in THP-1 or THP-1 HMGB1-Lucia cells was achieved using CRISPR-Cas9–mediated genome editing. For each target gene, a ribonucleoprotein (RNP) complex was assembled at a 1:1 molar ratio of single-guide RNA (sgRNA) and recombinant Cas9 (400 ng sgRNA and 2 µg Cas9) in Buffer R (Thermo Fisher Scientific), and incubated at room temperature for 15 min. A total of 0.6 × 10□ cells were harvested, washed with PBS (without calcium and magnesium), and incubated with the RNP complex. Electroporation was performed using the Neon NxT Electroporation System (Thermo Fisher Scientific) with the following parameters: 1700 V, 20 ns, and 1 pulse. Immediately after electroporation, cells were transferred into 24-well plates containing 500 µL of complete culture medium without antibiotics and incubated at 37°C with 5% CO_2_. After 4 days, single-cell sorting was performed into flat-bottom 96-well plates (one cell per well). Following 10 days of expansion, gene editing in individual clones was validated by PCR amplification and confirmed by Sanger sequencing. The sgRNA sequences used were: Non-targeting sgRNA #A35526, GSDMD 5’ ACTTCTACGATGCCATGGAT, HOIP 5’ GCGATTATATGGCTACACAG 3’, HOIL-1 5’ GAGACGCCACTGTCATATCA 3’, OTULIN 5’ TCAGGAGCTACGCTTAATCT 3’.

RNA extraction was performed on biological triplicates using the NucleoSpin RNA Plus kit (Macherey-Nagel). Purified RNA was reverse transcribed into cDNA using the Maxima First Strand cDNA Synthesis Kit (Thermo Fisher Scientific). Quantitative PCR (qPCR) was carried out using 25 ng of cDNA and PerfeCTa SYBR Green SuperMix Low ROX (QuantaBio). Gene expression levels were analyzed using the 2^−^ΔΔCt method and normalized to the housekeeping genes ACTB and HPRT1.The following primers were used:

ACTB forward, 5□-GGACTTCGAGCAAGAGATGG-3□; ACTB reverse, 5□-AGCACTGTGTTGGCGTACAG-3□; HPRT1 forward; 5□-TGACACTGGCAAAACAATGCA-3□; HPRT1 reverse, 5□-GGTCCTTTTCACCAGCAAGCT-3□; HOIP forward 5’-GAGCCCCGAAACTACCTCAAC-3’; HOIP reverse 5’-CTTGACACCACGCCAGTACC-3’; HOIL-1 forward 5’-TGCTCAGATGCACACCGTC-3’; HOIL-1 reverse 5’-CAAGACTGGTGGGAAGCCATA-3’; OTULIN forward 5’-GGGGCATCAGAACCGAGATTA-3’; OTULIN reverse 5’-TCGCCGTATGGAGGTGAACT-3’.

### Optogenetic model

THP-1 opto-ASC cells from ^29^ were washed twice with Opti-MEM, and 1-2 × 10□ cells were used per condition. Light-induced inflammasome activation was performed using a LED-based device (LAD-1 Driver, Amuva) placed inside the incubator directly beneath the multiwell plate containing the cells. Stimulation was carried out with blue light (477 nm) at 7.5 V, delivering 25 pulses per cycle, each pulse lasting 1 second. The stimulation cycle was repeated three times, with a one-minute interval between cycles. Following light exposure, cells were incubated for an additional 15-30 minutes to allow cell death to progress before collection for further analysis.

## REFERENCES

1. Newton, K., Strasser, A., Kayagaki, N., and Dixit, V.M. (2024). Cell death. Cell 187, 235– 256. 10.1016/j.cell.2023.11.044.

2. Bauernfeind, F.G., Horvath, G., Stutz, A., Alnemri, E.S., MacDonald, K., Speert, D., Fernandes-Alnemri, T., Wu, J., Monks, B.G., Fitzgerald, K.A., et al. (2009). Cutting Edge: NF-κB Activating Pattern Recognition and Cytokine Receptors License NLRP3 Inflammasome Activation by Regulating NLRP3 Expression. J. Immunol. 183, 787–791. 10.4049/jimmunol.0901363.

3. Zheng, D., Liwinski, T., and Elinav, E. (2020). Inflammasome activation and regulation: toward a better understanding of complex mechanisms. Cell Discov. 6, 36. 10.1038/s41421-020-0167-x.

4. Muñoz-Planillo, R., Kuffa, P., Martínez-Colón, G., Smith, B.L., Rajendiran, T.M., and Núñez, G. (2013). K+ Efflux Is the Common Trigger of NLRP3 Inflammasome Activation by Bacterial Toxins and Particulate Matter. Immunity 38, 1142–1153. 10.1016/j.immuni.2013.05.016.

5. Franklin, B.S., Bossaller, L., De Nardo, D., Ratter, J.M., Stutz, A., Engels, G., Brenker, C., Nordhoff, M., Mirandola, S.R., Al-Amoudi, A., et al. (2014). The adaptor ASC has extracellular and “prionoid” activities that propagate inflammation. Nat. Immunol. 15, 727–737. 10.1038/ni.2913.

6. Srinivasula, S.M., Poyet, J.-L., Razmara, M., Datta, P., Zhang, Z., and Alnemri, E.S. (2002). The PYRIN-CARD Protein ASC Is an Activating Adaptor for Caspase-1. J. Biol. Chem. 277, 21119–21122. 10.1074/jbc.C200179200.

7. Fernandes-Alnemri, T., Wu, J., Yu, J.-W., Datta, P., Miller, B., Jankowski, W., Rosenberg, S., Zhang, J., and Alnemri, E.S. (2007). The pyroptosome: a supramolecular assembly of ASC dimers mediating inflammatory cell death via caspase-1 activation. Cell Death Differ. 14, 1590–1604. 10.1038/sj.cdd.4402194.

8. Xia, S., Zhang, Z., Magupalli, V.G., Pablo, J.L., Dong, Y., Vora, S.M., Wang, L., Fu, T.-M., Jacobson, M.P., Greka, A., et al. (2021). Gasdermin D pore structure reveals preferential release of mature interleukin-1. Nature 593, 607–611. 10.1038/s41586-021-03478-3.

9. Rojas, E.R., Billings, G., Odermatt, P.D., Auer, G.K., Zhu, L., Miguel, A., Chang, F., Weibel, D.B., Theriot, J.A., and Huang, K.C. (2018). The outer membrane is an essential load-bearing element in Gram-negative bacteria. Nature 559, 617–621. 10.1038/s41586-018-0344-3.

10. Kayagaki, N., Kornfeld, O.S., Lee, B.L., Stowe, I.B., O’Rourke, K., Li, Q., Sandoval, W., Yan, D., Kang, J., Xu, M., et al. (2021). NINJ1 mediates plasma membrane rupture during lytic cell death. Nature 591, 131–136. 10.1038/s41586-021-03218-7.

11. Py, B.F., Kim, M.-S., Vakifahmetoglu-Norberg, H., and Yuan, J. (2013). Deubiquitination of NLRP3 by BRCC3 Critically Regulates Inflammasome Activity. Mol. Cell 49, 331–338. 10.1016/j.molcel.2012.11.009.

12. Song, N., Liu, Z.-S., Xue, W., Bai, Z.-F., Wang, Q.-Y., Dai, J., Liu, X., Huang, Y.-J., Cai, H., Zhan, X.-Y., et al. (2017). NLRP3 Phosphorylation Is an Essential Priming Event for Inflammasome Activation. Mol. Cell 68, 185-197.e6. 10.1016/j.molcel.2017.08.017.

13. Swatek, K.N., and Komander, D. (2016). Ubiquitin modifications. Cell Res. 26, 399–422. 10.1038/cr.2016.39.

14. Komander, D., Reyes-Turcu, F., Licchesi, J.D.F., Odenwaelder, P., Wilkinson, K.D., and Barford, D. (2009). Molecular discrimination of structurally equivalent Lys 63-linked and linear polyubiquitin chains. EMBO Rep. 10, 466–473. 10.1038/embor.2009.55.

15. Keusekotten, K., Elliott, P.R., Glockner, L., Fiil, B.K., Damgaard, R.B., Kulathu, Y., Wauer, T., Hospenthal, M.K., Gyrd-Hansen, M., Krappmann, D., et al. (2013). OTULIN Antagonizes LUBAC Signaling by Specifically Hydrolyzing Met1-Linked Polyubiquitin. Cell 153, 1312–1326. 10.1016/j.cell.2013.05.014.

16. Yang, X., Li, W., Zhang, S., Wu, D., Jiang, X., Tan, R., Niu, X., Wang, Q., Wu, X., Liu, Z., et al. (2020). PLK 4 deubiquitination by Spata2-CYLD suppresses NEK7-mediated NLRP3 inflammasome activation at the centrosome. EMBO J. 39, e102201. 10.15252/embj.2019102201.

17. Macé, Y.M., Bidère, N., and Douanne, T. (2025). The good, the bad, and the modified: CYLD’s post-translational tale. Front. Cell Death 4, 1583221. 10.3389/fceld.2025.1583221.

18. Iwai, K. (2021). LUBAC-mediated linear ubiquitination: a crucial regulator of immune signaling. Proc. Jpn. Acad. Ser. B 97, 120–133. 10.2183/pjab.97.007.

19. Douglas, T., Champagne, C., Morizot, A., Lapointe, J.-M., and Saleh, M. (2015). The Inflammatory Caspases-1 and −11 Mediate the Pathogenesis of Dermatitis in Sharpin-Deficient Mice. J. Immunol. 195, 2365–2373. 10.4049/jimmunol.1500542.

20. Rickard, J.A., Anderton, H., Etemadi, N., Nachbur, U., Darding, M., Peltzer, N., Lalaoui, N., Lawlor, K.E., Vanyai, H., Hall, C., et al. (2014). TNFR1-dependent cell death drives inflammation in Sharpin-deficient mice. eLife 3, e03464. 10.7554/eLife.03464.

21. Gurung, P., Sharma, B.R., and Kanneganti, T.-D. (2016). Distinct role of IL-1β in instigating disease in Sharpincpdm mice. Sci. Rep. 6, 36634. 10.1038/srep36634.

22. Nastase, M.-V., Zeng-Brouwers, J., Frey, H., Hsieh, L.T.-H., Poluzzi, C., Beckmann, J., Schroeder, N., Pfeilschifter, J., Lopez-Mosqueda, J., Mersmann, J., et al. (2016). An Essential Role for SHARPIN in the Regulation of Caspase 1 Activity in Sepsis. Am. J. Pathol. 186, 1206–1220. 10.1016/j.ajpath.2015.12.026.

23. Rodgers, M.A., Bowman, J.W., Fujita, H., Orazio, N., Shi, M., Liang, Q., Amatya, R., Kelly, T.J., Iwai, K., Ting, J., et al. (2014). The linear ubiquitin assembly complex (LUBAC) is essential for NLRP3 inflammasome activation. J. Exp. Med. 211, 1333–1347. 10.1084/jem.20132486.

24. Yu, Y., Yu, S., Lu, Z., Qiang, L., Zhong, Y., Ge, P., Lei, Z., Qiu, C., Fang, Y., Zhang, X., et al. (2025). Pathogenic phosphorylation of linear ubiquitin machinery causes inflammasome sensor degradation. Cell Rep. 44, 116286. 10.1016/j.celrep.2025.116286.

25. Katsuya, K., Oikawa, D., Iio, K., Obika, S., Hori, Y., Urashima, T., Ayukawa, K., and Tokunaga, F. (2019). Small-molecule inhibitors of linear ubiquitin chain assembly complex (LUBAC), HOIPINs, suppress NF-κB signaling. Biochem. Biophys. Res. Commun. 509, 700–706. 10.1016/j.bbrc.2018.12.164.

26. Heger, K., Wickliffe, K.E., Ndoja, A., Zhang, J., Murthy, A., Dugger, D.L., Maltzman, A., De Sousa E Melo, F., Hung, J., Zeng, Y., et al. (2018). OTULIN limits cell death and inflammation by deubiquitinating LUBAC. Nature 559, 120–124. 10.1038/s41586-018-0256-2.

27. Fiil, B.K., Damgaard, R.B., Wagner, S.A., Keusekotten, K., Fritsch, M., Bekker-Jensen, S., Mailand, N., Choudhary, C., Komander, D., and Gyrd-Hansen, M. (2013). OTULIN Restricts Met1-Linked Ubiquitination to Control Innate Immune Signaling. Mol. Cell 50, 818–830. 10.1016/j.molcel.2013.06.004.

28. Qiao, Y., Wang, P., Qi, J., Zhang, L., and Gao, C. (2012). TLR-induced NF-κB activation regulates NLRP3 expression in murine macrophages. FEBS Lett. 586, 1022–1026. 10.1016/j.febslet.2012.02.045.

29. Nadjar, J., Monnier, S., Bastien, E., Huber, A.-L., Oddou, C., Bardoulet, L., Leloup, H.B., Ichim, G., Vanbelle, C., Py, B.F., et al. (2024). Optogenetically controlled inflammasome activation demonstrates two phases of cell swelling during pyroptosis. Sci. Signal. 17, eabn8003. 10.1126/scisignal.abn8003.

30. Hu, J.J., Liu, X., Xia, S., Zhang, Z., Zhang, Y., Zhao, J., Ruan, J., Luo, X., Lou, X., Bai, Y., et al. (2020). FDA-approved disulfiram inhibits pyroptosis by blocking gasdermin D pore formation. Nat. Immunol. 21, 736–745. 10.1038/s41590-020-0669-6.

31. Haakonsen, D.L., and Rape, M. (2019). Branching Out: Improved Signaling by Heterotypic Ubiquitin Chains. Trends Cell Biol. 29, 704–716. 10.1016/j.tcb.2019.06.003.

